# Chromap Suite: an open-source single-binary platform for agentic multiomic RNA + ATAC profiling

**DOI:** 10.64898/2026.06.02.729736

**Authors:** Ling-Hong Hung, Ka Yee Yeung

## Abstract

**Background:** Single-cell multiomic profiling of RNA expression and chromatin accessibility is now a standard tool for resolving regulatory state in single cells, but existing analysis toolchains have lagged. Cell Ranger ARC, the proprietary multiomic pipeline, uses a custom broad peak caller rather than the MACS3 narrow peaks that the ATAC field has consolidated on, and its restrictive end-user licence forbids redistribution of analysis pipelines that include it. A fully open-source, permissively-licensed alternative anchored on community-standard methods—Chromap for ATAC alignment and MACS3 for narrow peak calling—has been impractical to assemble because the two codebases are written in different languages with incompatible runtimes, leaving practitioners to chain them together with ad-hoc scripts.

**Results:** We present Chromap Suite, the chromatin-accessibility side of an open-source multiomic stack built in support of the NIH Molecular Phenotypes of Null Alleles in Cells (MorPhiC) consortium’s multiomic production pipeline. We extended Chromap with native BAM output and coordinate sorting, in-process narrow peak calling, optional Y-chromosome filtering, and native input from the compressed binary CBQ sequencing format alongside FASTQ, and hardened the result with a regression-test matrix that auto-validates the four upstream Chromap presets (bulk ATAC, scATAC, ChIP-seq, Hi-C). We reimplemented MACS3’s narrow peak caller in portable C++ as `libMACS3`, byte-identical to MACS3 v3.0.3 and free of any Python interpreter dependency. Finally, we extracted Chromap’s alignment and fragment-generation paths into a callable C++ library (`libchromap`) and embedded both `libchromap` and `libMACS3` into STAR Suite, so that one `STAR` invocation runs alignment, peak calling, and cell calling for both RNA and ATAC modalities concurrently—to our knowledge the first true single-binary RNA + ATAC multiomic implementation. On the public 3K PBMC Multiome at 32 threads, the platform completes in 18 minutes 55 seconds wall time and 44.6 GB peak resident memory, against 40 minutes 4 seconds and 79.1 GB resident memory for Cell Ranger ARC v2.2.0—a 2.12 × wall speedup with 1.8× less peak memory—and produces 50,274 peaks that are byte-identical to MACS3 v3.0.3. To support deployment by both research scientists and the AI agents increasingly used in bioinformatics analysis, Chromap Suite ships a Model Context Protocol (MCP) server and a browser-based Launchpad driven by a shared set of composable YAML recipes that humans and agents drive the same way.

**Conclusions:** Chromap Suite delivers a unified, freely redistributable multiomic pipeline that produces the MACS3 narrow peaks downstream ATAC analyses already rely on, with substantially lower wall time and memory than the proprietary alternative. The MIT- and BSD-3-licensed code carries no redistribution restrictions, the constituent libraries are independently embeddable in other open-source tools, and the MCP server plus Launchpad recipes make the platform straightforward to drive both by humans and by AI agents.

## 1 Background

Single-cell multiomic profiling of RNA expression and chromatin accessibility has become a standard tool for resolving regulatory state by providing independent corroborative data in the same cell. However the development of existing toolchains for processing these datasets has lagged behind. For transcriptomics-derived workflows (e.g., Perturb-seq), the proprietary Cell Ranger software anchors a stable, standard-methods stack. However, its multiomic counterpart Cell Ranger ARC [2], uses its own custom ATAC broad peak caller rather than the standard MACS3 [15] caller that produces narrow better resolved peaks. As a result, many groups routinely re-call ATAC peaks with MACS3 when using ARC. Cell Ranger variants are also restrictively licensed with a EULA that forbids redistribution of analysis pipelines that include it which rules out a shareable analysis stack using ARC as an anchor.

An effective open-source alternative has been difficult to develop. Cell Ranger does use STAR [5] internally and STAR Solo was developed as a scRNA-seq alternative. However, STAR Solo [8] was designed to mimic early Cell Ranger releases and has diverged from current versions. The new STAR Suite [7] (https://github.com/morphic-bio/STAR-suite) extends STAR with integrated C++ modules for rapid and scalable analysis of bulk and single cell transcriptomics experiments and has higher parity with the current Cell Ranger. On the ATAC-seq side, Chromap [13] has emerged as the modern aligner and fragment caller and MACS3, the preferred peak caller. Chromap is written in C++ and MACS3 in Python with Cython libraries for speed. The complexity of the codebases for both these packages has made it a practical impossibility for third parties to integrate the two. This has changed with the advent of AI-assisted software engineering techniques, and we were able to integrate Chromap and MACS3 into Chromap Suite and then integrate Chromap Suite into STAR Suite using the same human-directed agentic development methods that we used previously to create STAR Suite.

In support of the NIH Molecular Phenotypes of Null Alleles in Cells (MorPhiC) consortium’s multiomic production pipeline, we extended and hardened Chromap to deliver the downstream stages of a complete ATAC pipeline that the standalone tool leaves to external orchestrators. Chromap gained native BAM output and coordinate sorting (replacing the conventional `chromap` — `samtools sort` — `samtools index` chain), in-process narrow peak calling, optional Y-chromosome filtering, native input from the ARC Institute’s compressed binary CBQ sequencing format [3] alongside FASTQ, and a regression-test matrix that auto-validates the four upstream Chromap presets (bulk ATAC, scATAC, ChIP-seq, Hi-C). To make narrow peak calling embeddable inside another C++ tool without bringing in a Python interpreter, MACS3’s narrow-peak path was reimplemented in portable C++ as `libMACS3`, byte-identical to MACS3 v3.0.3. Finally, we integrated these pieces into a single open-source binary: Chromap’s alignment and fragment-generation paths were extracted into a callable C++ library (`libchromap`) and embedded together with `libMACS3` into STAR Suite, so that one `STAR` invocation runs alignment, peak calling, and cell calling for both RNA and ATAC modalities. To our knowledge this is the first true single-binary RNA + ATAC multiomic implementation, and it is substantially faster and lighter on memory than the proprietary alternative (`Multiomic RNA + ATAC pipeline` under Results, Section 2.3). Both `libchromap` and `libMACS3` are also distributed as standalone static C++ archives that other open-source tools can embed directly: `libchromap` retains Chromap’s lean native dependencies (zlib, pthread) and bundles htslib for BAM read/write/sort; `libMACS3` depends only on htslib (used solely for the Poisson tail in `kfunc`) and zlib (for gzipped fragment input). Neither library pulls in a Python/Cython runtime—both build to small static archives that drop into a host process without an interpreter.

The main design goal of Chromap Suite, beyond the speed gains above, was to simplify deployment of chromatin-accessibility and multiomic workflows for both research scientists and AI agents. Agent-assisted analysis is becoming central to bioinformatics, and the open-source, permissively-licensed (MIT and BSD-3) nature of Chromap Suite removes the redistribution barriers that block agent-driven use of ARC-anchored stacks. Modern agentic environments increasingly rely on *skills*—structured, declarative workflow definitions (a name, description, parameter slots, and optional preflight directives) that an agent can invoke as named units rather than improvising one from scratch using an expensive LLM. Chromap Suite ships a simplified instance of this pattern: each task (indexing, bulk ATAC, scATAC, multiomic, ChIP-seq, Hi-C) is captured as a small YAML recipe declaring inputs, outputs, reasonable defaults, and preflight checks. The MCP server—built on the Model Context Protocol, a standard agent tool-server protocol—and the browser-based Launchpad are two faces of the same underlying server: the MCP face exposes the recipes to agents as schema-validated tool calls, and the Launchpad face renders the same recipes as web forms for human users. Two factors improve robustness, lower token use and cost: the unified-binary architecture reduces the number of tool boundaries an agent must traverse, and serving recipes through the MCP server makes these tools easily discoverable and usable by agents.

## 2 Results

### 2.1 Bulk ATAC-seq pipeline

`libchromap` inherits Chromap’s bulk-ATAC alignment capability without modification: bulk and single-cell ATAC inputs share the same library and CLI, and the libMACS3 validation and runtime numbers reported in the Methods (`libMACS3`, Section 5.2) were measured on the 3K PBMC ATAC reads—essentially a bulk-ATAC measurement aside from the per-cell barcode-assignment step that distinguishes the single-cell regime. The peak-calling code path is identical between the two regimes, so the byte-identical-MACS3 parity and the wall-clock and thread-scaling numbers (Fig. 1) carry over to bulk-ATAC inputs directly; the downstream output simply replaces per-cell calls with a single bulk per-sample peak file.

**Figure 1:**
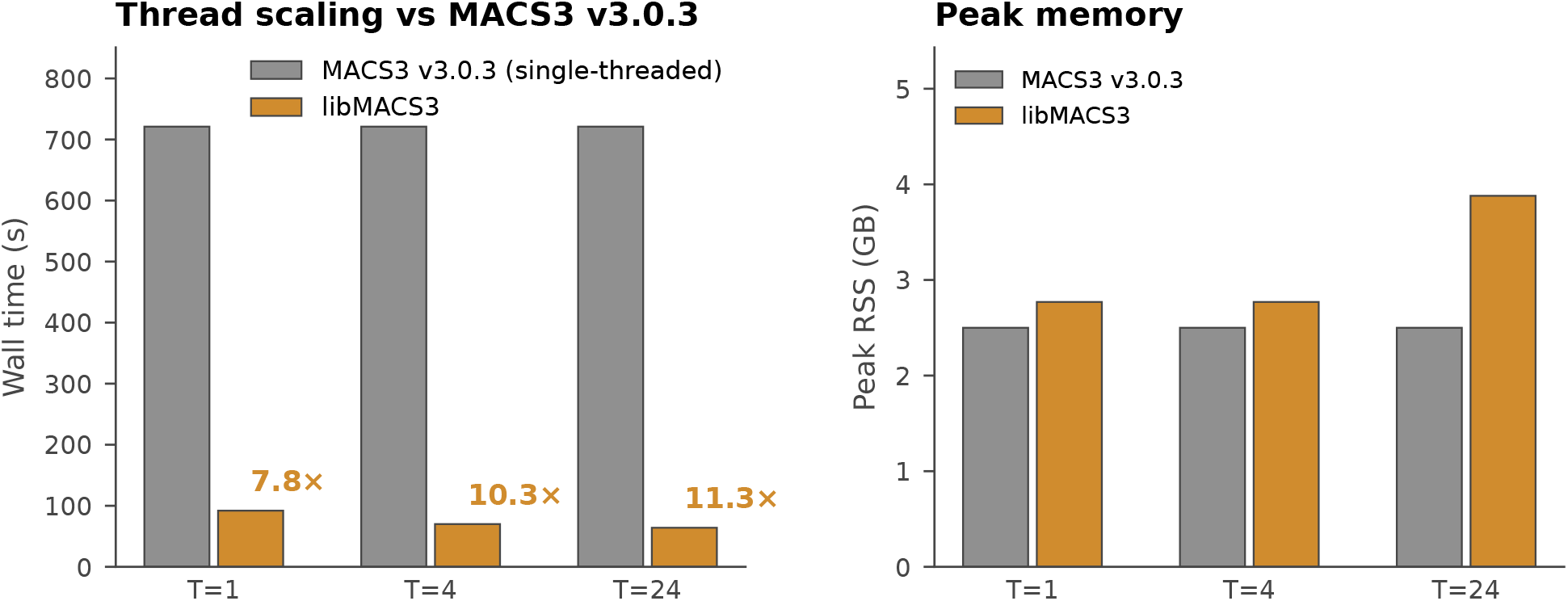
libMACS3 runs 7.8–11.3× faster than Cython MACS3 v3.0.3 with byte-identical output. Standalone wall-clock (left) and peak resident memory (right) at *T* = 1, 4, 24 threads versus single-threaded MACS3 v3.0.3 on the public 10x 3K PBMC Multiome ATAC channel (53.97 M deduplicated fragments). Speedup ratios annotated above libMACS3 bars. All thread counts produce a 50,274-peak narrowPeak file byte-identical to the MACS3 v3.0.3 reference; treat, lambda, and `ppois` bedGraphs and summit calls are likewise byte-identical (see Methods, Section 5.2).

### 2.2 Single-cell ATAC-seq pipeline

The 3K PBMC Multiome ATAC channel doubles as a standalone single-cell ATAC benchmark: the same input that anchored libMACS3 validation (Section 5.2) produces 50,274 byte-identical narrowPeak peaks via `libchromap` and `libMACS3` in 1 min 32 s wall time at single-thread (Fig. 1). The integrated path delivers the MACS3-equivalent narrow peaks scATAC analyses already rely on through a single binary invocation, replacing the conventional chained `chromap` + MACS3 toolchain that scATAC pipelines—e.g., SnapATAC2 [14] and ArchR [6], and the Signac [12] R package for Seurat-based multiome workflows—typically wrap around.

### 2.3 Multiomic RNA + ATAC-seq pipeline

Joint RNA + ATAC multiomic processing is the use case where the integration value of `libchromap` and `libMACS3` pays off most clearly. Instead of running STAR Suite [7], Chromap, MACS3, and an ATAC cell caller as a chained pipeline coordinated by Nextflow or Snakemake, the entire workflow— alignment, GEX UMI counting, GEX cell calling, ATAC mapping, narrow peak calling, ATAC peak/MEX materialization, and ATAC per-barcode cell calling—runs inside a single `STAR` invocation. STAR’s GEX path and `libchromap`’s ATAC path execute concurrently under a shared permit allocator (Section 5.1), emitting sorted BAM, the standard fragments TSV, and a binary ATAC fragment sidecar. When the concurrent mapping phase completes and chromap’s mapping state is freed, STAR invokes `RunMultiomeAtacPeakMex` (in `core/features/libchromap_contract/`) as an in-process post-alignment phase: the same library reads the sidecar from disk and produces MACS3-equivalent narrow peaks via `libMACS3` (Section 5.2) together with the ATAC peak MEX, ATAC per-barcode cell call (via libscrna’s empty-cells function), and ATAC metrics. The sidecar boundary is internal to the STAR process but kept on disk rather than in RAM so the materialization phase runs at much lower peak resident memory than the alignment phase. The same `RunMultiomeAtacPeakMex` library is also exposed as a standalone CLI binary (`star_multiome_atac_peak_mex`) for re-running just the materialization step against an existing sidecar—useful for debugging and for incremental re-runs without re-aligning.

#### Cross-modal cell calling

Two cell-calling components run in-process. GEX cell calling uses STAR Suite’s `libscrna` library, a refactored `EmptyDrops_CR` implementation that produces Cell Ranger-compatible cell calls and is distinct from the older version embedded in STAR Solo. For ATAC, libchromap emits an *ATAC fragment sidecar* (`AEV1` format: 32-byte header followed by 24-byte fragment records keyed by 16-byte barcode, with a `(size−32)/24` record-count parity check) alongside its fragment output during alignment; libscrna’s empty-cells function consumes the sidecar together with the libMACS3 peak output to compute per-barcode evidence and call cells. The sidecar separates fragment production from cell-call-time aggregation, decouples the two phases for independent verification, and keeps the entire pipeline inside the single `STAR` invocation with no external orchestration. On the headline run the sidecar contains 53,969,811 records.

#### End-to-end benchmark on 3K PBMC (Fig. 4)

On the public 10x 3K PBMC Multiome dataset [1] at 32 threads, the integrated pipeline produces sorted-and-indexed BAM, fragments, narrowPeak, summits, the ATAC fragment sidecar, the GEX expression matrix, ATAC peak MEX (50,274 × 315,435 with 15.31 M nonzeros), ATAC metrics, and ATAC + GEX cell calls in 18 min 55 s wall time (1134.81 s), with a peak resident memory of 44.55 GB measured via `/usr/bin/time -v` on the single `STAR` process. The inline post-alignment materialization phase contributes ~83 s to the wall (STAR-log timestamps). The same input under Cell Ranger ARC v2.2.0 (`cellranger-arc count` `--create-bam=true --nosecondary --disable-cell-annotation --localcores=32`) takes 40 min 4 s and 79.07 GB peak RSS—a 2.12× wall speedup and a 1.8× less peak memory. Output hashes (narrowPeak, summits, ATAC metrics, peak MEX, GEX raw/filtered matrices) are byte-identical between the inline-materialization run and an earlier two-step run, confirming the architectural collapse to a single `STAR` invocation preserves results. Scope is matched: the same set of stages ARC bundles inline (alignment, GEX UMI counting, GEX `EmptyDrops_CR`, ATAC mapping, peak calling, ATAC per-barcode cell call). ARC’s RSS is read from cellranger’s own `_perf` JSON because the parent-process `/usr/bin/time -v` reading is misleading—ARC forks 252 child processes at peak; the integrated pipeline’s RSS reading is single-process and accurate. GEX `nCellsSimple` on this run was 2,974 (`Gene`) and 2,976 (`GeneFull`).

#### Peak parity with MACS3

The narrowPeak file produced by the integrated pipeline is byte-identical to the output of standalone MACS3 v3.0.3 on the same input (50,274 peaks), and the summits file is similarly byte-identical. Cell Ranger ARC produces 81,157 peaks via its own custom peak caller (stages `_PEAK_CALLER`, `COUNT_CUT_SITES`, `DETECT_PEAKS`, `CONVERT_SIGNAL_TRACK`), which is not a parameterisation of MACS—peak counts and boundaries between the two outputs are not directly comparable. ARC users routinely re-call peaks with MACS for downstream analysis; that re-calling step is unnecessary with our pipeline.

#### Allocator behaviour on this run

STAR Suite’s permit allocator (Section 5.1) coordinated the concurrent GEX and ATAC threads. Its goal is to share the thread budget between the two domains so that they finish at approximately the same time, since whichever domain finishes earlier then sits idle while the other completes alone. We evaluated several thread-distribution strategies during development; the one used here was the one that consistently produced the shortest wall time across our test scenarios on this hardware, and is what makes the 18-minute wall reproducible.

## 3 Discussion

In support of the MorPhiC consortium’s multiomic production pipeline, we extended and hardened Chromap’s ATAC path: native BAM output with coordinate sorting and indexing replaces the conventional samtools chain, in-process narrow peak calling avoids the intermediate fragments-file write, native input from the compressed binary CBQ sequencing format complements FASTQ, and a four-preset regression matrix guards the existing Chromap behaviour against regression. The MACS3 narrow peak caller was reimplemented in portable C++ as `libMACS3`, byte-identical to MACS3 v3.0.3, so peak calling runs inside the same process as alignment without an embedded Python interpreter. Finally, we integrated these pieces into a single open-source binary: by extracting Chromap’s alignment path into a callable C++ library (`libchromap`) and embedding both into STAR Suite, one `STAR` invocation now runs alignment, peak calling, and cell calling for both RNA and ATAC modalities concurrently. To our knowledge this is the first true single-binary RNA + ATAC multiomic implementation, and the whole stack is freely redistributable under permissive (MIT and BSD-3) licences. Chromap and MACS3 already represent the state-of-the-art open-source ATAC toolchain that downstream pipelines such as SnapATAC2 [14], ArchR [6], and Signac [12] consume; what has been missing is an integrated, redistributable open-source pipeline that combines them in a single tool. Cell Ranger ARC instead uses a custom wide peak caller that departs from the MACS3 narrow-peak standard the ATAC field has otherwise consolidated on, and its license forbids redistribution of analysis pipelines that include it.

There is also an architectural difference between the two pipelines. Cell Ranger ARC is implemented as an internal workflow of many Python stages that pass intermediate results to one another through files on disk—252 child processes are forked at peak on the 3K PBMC benchmark (Section 2.3). Chromap Suite runs the equivalent set of stages inside a single binary, sharing memory directly between alignment, peak calling, and cell calling with no inter-stage file handoffs. The MorPhiC consortium and other users can still wrap Chromap Suite in higher-level orchestration scripts—this is what the companion `morphic-recipes` repository (Section 5.1) does for production deployment—but the orchestration sits *outside* the binary rather than inside it. This is one source of the lower peak resident memory reported below, and is what lets the ATAC pipeline embed cleanly inside another single-process tool (STAR Suite, for the multiomic case).

The open-source pipeline is also faster and more memory-efficient than the proprietary alternative. The integrated multiomic workflow runs 2.12× faster than Cell Ranger ARC at matched scope on the public 3K PBMC Multiome benchmark, with 1.8× less peak resident memory (Section 2.3). libMACS3 on its own runs 7.8–11.3× faster than Cython MACS3 across one to twenty-four threads (Section 5.2).

A growing share of bioinformatics analyses is now driven by AI agents rather than only by human users. A workflow stitched together from several command-line tools is awkward for an agent to drive: at every step the agent has to find the right binary, parse an intermediate file, and recover from format mismatches, and any one of those is a place a small mistake can derail the run. Putting all of the steps inside one binary with one set of arguments and one named set of outputs eliminates most of these sources of failure, and the workflow is then simple enough that smaller in-house models—not only the largest frontier-grade ones—can drive it reliably. Chromap Suite is designed for this: an MCP server makes the workflows available to agents as named, parameterised recipes; a browser-based Launchpad presents the same recipes as web forms for human users; and an `AGENTS.md` file in the repository describes the layout for agents that need to navigate it. The same recipe definitions back both the agent calls and the human forms, so what an agent runs and what a person reads are the same workflow.

Chromap Suite is the chromatin-accessibility companion to STAR Suite [7], and the two compose for the joint RNA + ATAC multiomic case. They share the same design philosophy—a single open-source binary that humans and AI agents can drive equally well—but their scopes differ because the underlying tools differ. STAR Suite is the larger effort because the proprietary alternative, Cell Ranger, has accumulated many capabilities that STAR by itself does not provide (perturb-seq, 10x Flex, SLAM-seq, gzip handling, adapter trimming, `EmptyDrops_CR` cell calling); STAR Suite adds all of them to reach parity. The ATAC field is structured differently. Chromap already provides adapter trimming, barcode correction, deduplication, and Tn5 shifting as built-in capabilities, so less had to be added to reach a complete pipeline. The additions we did make are nonetheless substantive (Tables 1, 2, 3): native BAM output with coordinate sorting (replacing the conventional `chromap` — `samtools sort` — `samtools index` chain); in-process narrow peak calling that hands fragments directly to libMACS3 inside the same process, with no intermediate file written to disk; a compact binary sidecar file that carries fragments to the ATAC cell caller; a new low-memory mode that supports any combination of these output choices; native input from the ARC Institute’s compressed binary CBQ sequencing format alongside FASTQ; and the `libchromap` library that lets a host process embed the whole pipeline.

**Table 1:**
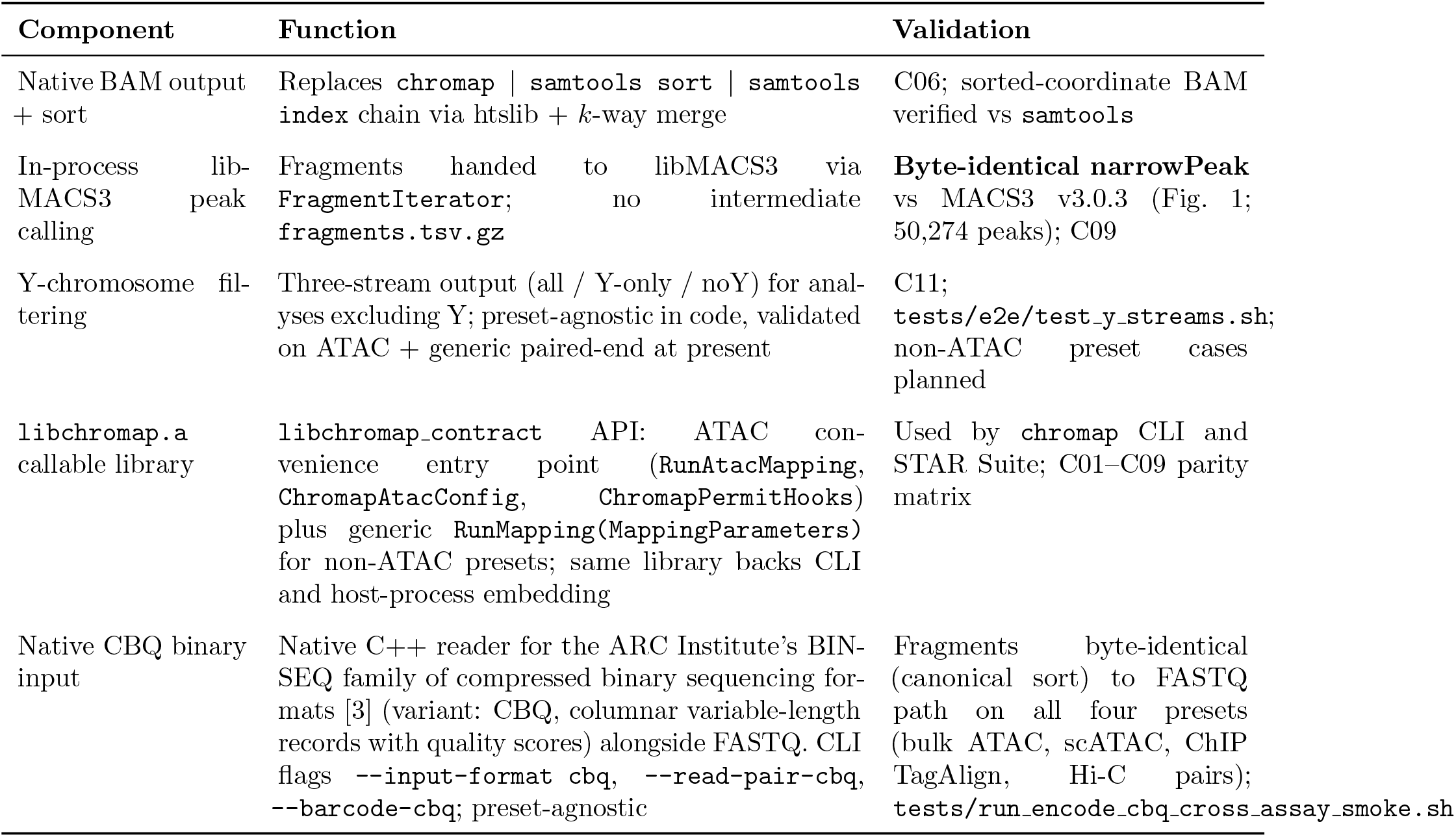
Chromap Suite additions to Chromap: core additions. Extensions to the standalone tool that bring Chromap to a complete pipeline (BAM output, in-process peak calling, callable library). Validation references denote regression-matrix cases (C01–C11) and parity figures. Companion tables: Table 2 (reliability and tooling) and Table 3 (ATAC-multiomic).

**Table 2:**
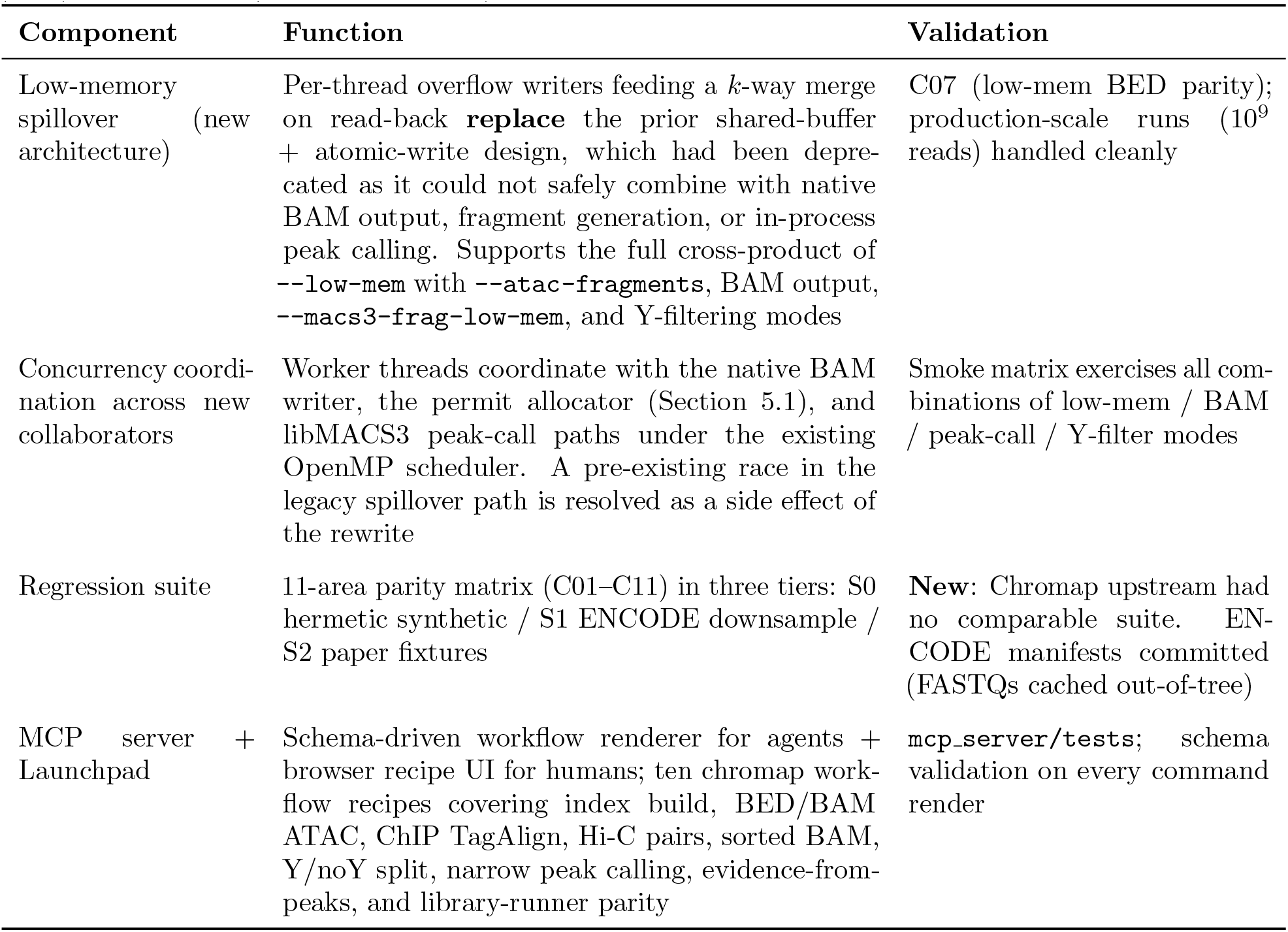
Chromap Suite additions to Chromap: reliability and tooling additions. Improvements to quality, reliability, and developer/agent accessibility. Companion tables: Table 1 (core) and Table 3 (ATAC-multiomic).

**Table 3:** Chromap Suite additions to Chromap: ATAC-multiomic additions. Components enabling composition with STAR Suite for joint RNA + ATAC processing in a single binary. Companion tables: Table 1 (core) and Table 2 (reliability and tooling).

| Component | Function | Validation |
| --- | --- | --- |
| ATAC fragment sidecar | Compact binary file emitted alongside the BAM and fragments TSV; one barcode-keyed record per fragment (24 bytes) lets libscrna’s empty-cells function call cells without re-parsing the gzipped fragments TSV | AEV1 magic header + (size-32)/24 == record-count parity check; 53,969,811 records on the 3K PBMC headline run |
| Multiomic integration with STAR Suite | libchromap runs as a STAR Suite worker thread (concurrent ATAC + GEX); after alignment, STAR invokes RunMultiomeAtacPeakMex in-process for narrow peaks, peak MEX, ATAC metrics, and ATAC cell calls—all inside a single STAR invocation | **18:55 / 44.6 GB / 2.12× vs ARC v2.2.0** (Fig. 4); MACS3-equivalent narrow peaks |

Our work focuses on the ATAC-seq path because that is the chromatin-accessibility data type the MorPhiC consortium processes at production scale, and Chromap Suite was built in service of that pipeline. The existing Chromap functionality is preserved unchanged: all upstream CLI flags, the assay-type presets (ATAC, ChIP-seq, Hi-C), and the legacy output formats continue to behave exactly as before, and the new additions are activated by new flags whose defaults match the upstream behaviour, so existing chromap scripts and pipelines continue to work without modification. Several of the new additions—notably native BAM sort/index and CBQ binary input—are not ATAC-specific in the code and apply to the other Chromap assay types as well, and Chromap Suite ships a regression matrix (Section 5.1) that exercises all four upstream presets (bulk ATAC, single-cell ATAC, ChIP-seq, and Hi-C) against both synthetic and real ENCODE data to confirm this. What remains ATAC-specific is the in-process libMACS3 narrow-peak integration: an analogous integration would be meaningful for ChIP-seq but not for Hi-C, and we have not yet built it because doing so cleanly needs a real downstream use case to anchor the parameter choices and the reference dataset, and the MorPhiC consortium does not currently process ChIP-seq at production scale. The underlying infrastructure to add such integrations when a use case arrives is already in place: the scheduler that coordinates concurrent ATAC and GEX work in this paper was generalised during this work so that further processing domains can be added without further redesign.

## 4 Conclusions

Chromap Suite is a unified, fully open-source multiomic chromatin-accessibility pipeline that produces the MACS3 narrow peaks downstream ATAC analyses already rely on, packaged as a single binary that runs alignment, peak calling, and cell calling for both ATAC and RNA modalities without any external workflow manager. The combination of `libchromap` (an embeddable form of Chromap with built-in BAM output and peak calling) and `libMACS3` (a portable C++ reimplementation of MACS3’s narrow peak caller) closes the gap left by proprietary multiomic platforms that depart from community-standard methods, and the permissive (MIT and BSD-3) licensing means the whole platform can be freely redistributed as part of downstream analysis pipelines.

Beyond the speed and memory gains over the proprietary alternative, the platform is designed to be straightforward to drive both by humans and by AI agents. Workflows are exposed through an MCP server and a browser Launchpad as a small set of named recipes that an agent can call directly and that a person can fill in as a web form. Both audiences see the same recipe, and because the pipeline runs as a single binary, an agent does not have to bridge between separate tools mid-run—which lowers the per-run cost and the chances of a small mistake derailing the run. Chromap Suite is intended both as a practical pipeline for current scATAC and joint RNA + ATAC analyses and as a concrete example of how community-standard bioinformatics tools can be packaged as open-source libraries that humans and agents can drive equally well.

## 5 Methods

### 5.1 libchromap: extending Chromap for end-to-end use

Chromap [13] is a fast ATAC-seq aligner, but its standalone form leaves the downstream stages of a complete pipeline—coordinate-sorted BAM, BAM indexing, and narrow peak calling—to external tools chained through pipeline orchestrators. We extended Chromap to integrate those stages directly into the C++ codebase and extracted the result as `libchromap`, an embeddable callable library that exposes ATAC mapping, fragment generation, native BAM output, and in-process peak calling through a single C++ API (source: https://github.com/morphic-bio/Chromap-suite). Tables 1–3 summarise Chromap Suite’s additions, organised into core capabilities, reliability and tooling infrastructure, and ATAC-multiomic integration pieces; the remainder of this section describes each in detail.

#### Single-binary alignment-to-peaks (Fig. 2)

**Figure 2:**
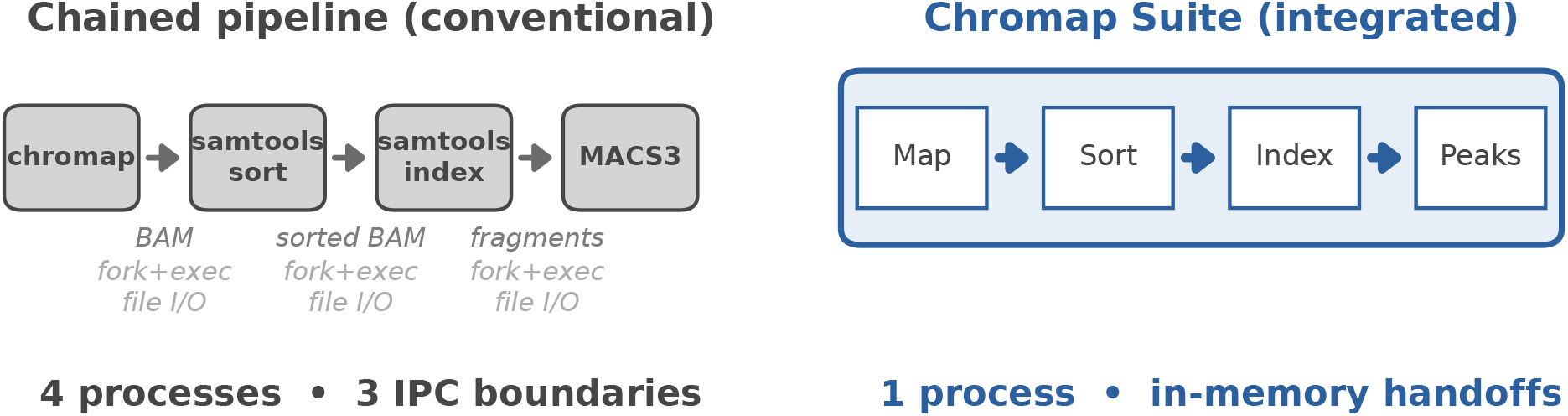
Chromap Suite collapses the conventional ATAC toolchain into a single in-process pipeline. The conventional four-process chained pipeline (`chromap` — `samtools sort` — `samtools index` — MACS3) is replaced by a single `libchromap` process whose internal sub-blocks (mapping, sort, index, libMACS3 peak call) communicate in-memory, eliminating three IPC boundaries.

`libchromap` writes BAM output natively via htslib [4] and performs a coordinate sort with *k*-way disk-merge spillover, eliminating the conventional `chromap` | `samtools sort` | `samtools index` chain [9] and the inter-process file write/read steps it requires. Fragment output is similarly first-class. Peak calling proceeds in-process: fragments are passed to `libMACS3` (Section 5.2) through the `FragmentIterator` API without an intermediate `fragments.tsv.gz` write, so a single `libchromap` invocation produces sorted-and-indexed BAM, fragments, and MACS3-equivalent narrowPeak and summit outputs in one process.

#### Native CBQ binary input

Chromap Suite reads the ARC Institute’s BINSEQ family of compressed binary sequencing formats [3] natively, in addition to FASTQ. The variant used is CBQ (columnar variable-length records), which carries quality scores, headers, and ambiguous bases and is therefore a lossless substitute for FASTQ. A native C++ CBQ reader (`src/cbq_reader.` {`h,cc`}, `src/cbq_batch_producer.` {`h,cc`}) decodes records directly into Chromap’s existing `SequenceBatch` buffers, so the reader hooks in below the preset dispatch and is preset-agnostic. CBQ inputs are driven by three CLI flags (`--input-format cbq`, `--read-pair-cbq`, `--barcode-cbq`) that mirror the FASTQ flags; the same flags compose with all assay presets and with the new BAM, fragments, and sidecar outputs. The integration removes the intermediate FASTQ write that a typical upstream pipeline emits before alignment, and lets Chromap Suite participate directly in pipelines that already store reads in CBQ form. CBQ inputs are validated on all four upstream presets by the regression matrix described below (`tests/run_encode_cbq_cross_assay_smoke.sh`); fragments are byte-identical (under canonical sort) to the FASTQ-input variant for bulk ATAC, single-cell ATAC, ChIP-seq TagAlign, and Hi-C pairs.

#### Library API for permit-shared embedding

`libchromap.a` exposes its capabilities through the `libchromap contract` boundary: an ATAC-focused convenience entry point `RunAtacMapping()`, a `ChromapAtacConfig` struct that bundles the reference FASTA, index path, FASTQ inputs, barcode whitelist, and output paths, and a `ChromapPermitHooks` struct that lets the caller participate in thread coordination. The library was extracted specifically to allow direct integration with STAR Suite through its permit allocator (described in the next paragraph) rather than running `chromap` as an external subprocess: a shared thread-coordination namespace lets the host process throttle ATAC and GEX work against a single permit budget, avoiding the unregulated oversubscription that occurs when the alternative subprocess approach is used. The same library backs the standalone `chromap` CLI binary unchanged.

#### Scope of the callable surface, and feasibility of extending it

This paragraph concerns the *library-API capability* for non-ATAC presets, which is distinct from the consortium-level multiomic ChIP+RNA and Hi-C+RNA integrations the Background section deferred. The convenience entry point `RunAtacMapping()` is one of two libchromap entry points; the other is the generic `RunMapping()` that accepts a fully populated `MappingParameters` struct (the same struct the CLI fills in for every preset). `RunMapping()` drives the same underlying paired-end mapping template that the CLI’s `--preset atac`, `--preset chip`, and `--preset hic` paths share; output dispatch over BED, TagAlign, PAIRS, SAM, BAM, and CRAM is already template-polymorphic in the library. A host process embedding libchromap can therefore already drive ChIP-seq or Hi-C alignment through `RunMapping()` without extending the binary, and the native BAM output and Y-chromosome filtering capabilities described above are implemented preset-agnostically and are expected to work for any preset. We have not yet added preset-specific convenience wrappers (`RunChipMapping()`, `RunHicMapping()`) or extended the regression matrix’s BAM and Y-filter cases to non-ATAC presets, because the MorPhiC consortium pipeline that drives this work consumes only ATAC data and we have not had a downstream caller asking for them. Adding the convenience wrappers is a small change—each is a thin parameter-setting delegation to `RunMapping()` —and rounding out the callable surface for ChIP-seq and Hi-C, with matching parity tests against the corresponding CLI invocations, is a natural follow-on. Building out a full single-binary ChIP-seq+RNA or Hi-C+RNA multiomic pipeline of the kind delivered here for ATAC remains a separate engineering effort (Discussion, Section 3) and is out of scope for the present work.

#### Concurrent execution alongside STAR (Fig. 3)

When embedded in STAR Suite via the contract, `libchromap` runs as a worker thread that begins ATAC mapping in parallel with STAR’s GEX mapping; the two compete for cycles on a shared thread budget. STAR Suite includes a *permit allocator* that partitions that budget between concurrent processing domains (GEX mapping, feature processing, ATAC mapping) and rebalances them toward finishing at approximately the same time. The target is not equitable CPU access—which the OS scheduler already provides—but efficient pipeline completion: whichever domain finishes earlier sits idle while the others complete, and that trailing serial tail is the dominant source of inefficiency in concurrent multi-domain pipelines. To support the multiomic case here, an ATAC domain was added to the allocator. We tried several thread-distribution strategies during development; the one shipped here was the one that consistently produced the shortest wall time across our test scenarios. A Gantt of a representative run on the public 10x 3K PBMC Multiome dataset (Fig. 3, drawn from STAR log timestamps) shows ATAC mapping completing well before STAR’s GEX path, so the integrated end-to-end wall is dominated by STAR’s GEX phase rather than by chromap or peak calling. The final *peak + expression matrix* step on the STAR Suite lane is a join point between the two sides: it depends on the cell barcodes produced by GEX cell calling and on the fragments plus AEV1 sidecar produced by the ATAC side, and is therefore run after both lanes complete.

**Figure 3:**
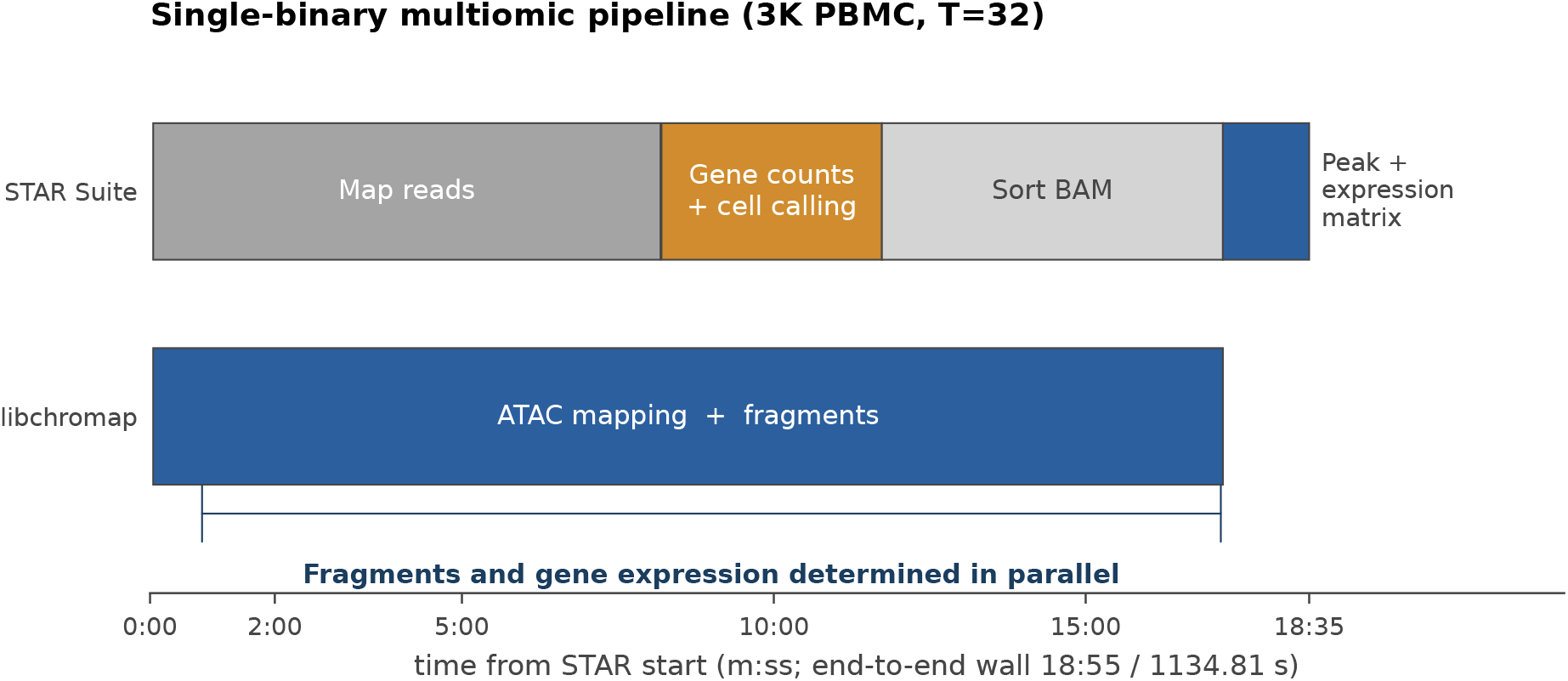
Concurrent-execution Gantt for the multiomic run on the public 3K PBMC Multiome at 32 threads (drawn from STAR-log timestamps). The fragments side (`libchromap`, bottom lane) and the gene-expression side (STAR, top lane) are determined in parallel under the permit allocator; the trailing *peak + expression matrix* step on the STAR Suite lane joins outputs from both sides (see Methods, Section 5.1). End-to-end wall: 18:55 (1134.81 s).

**Figure 4:**
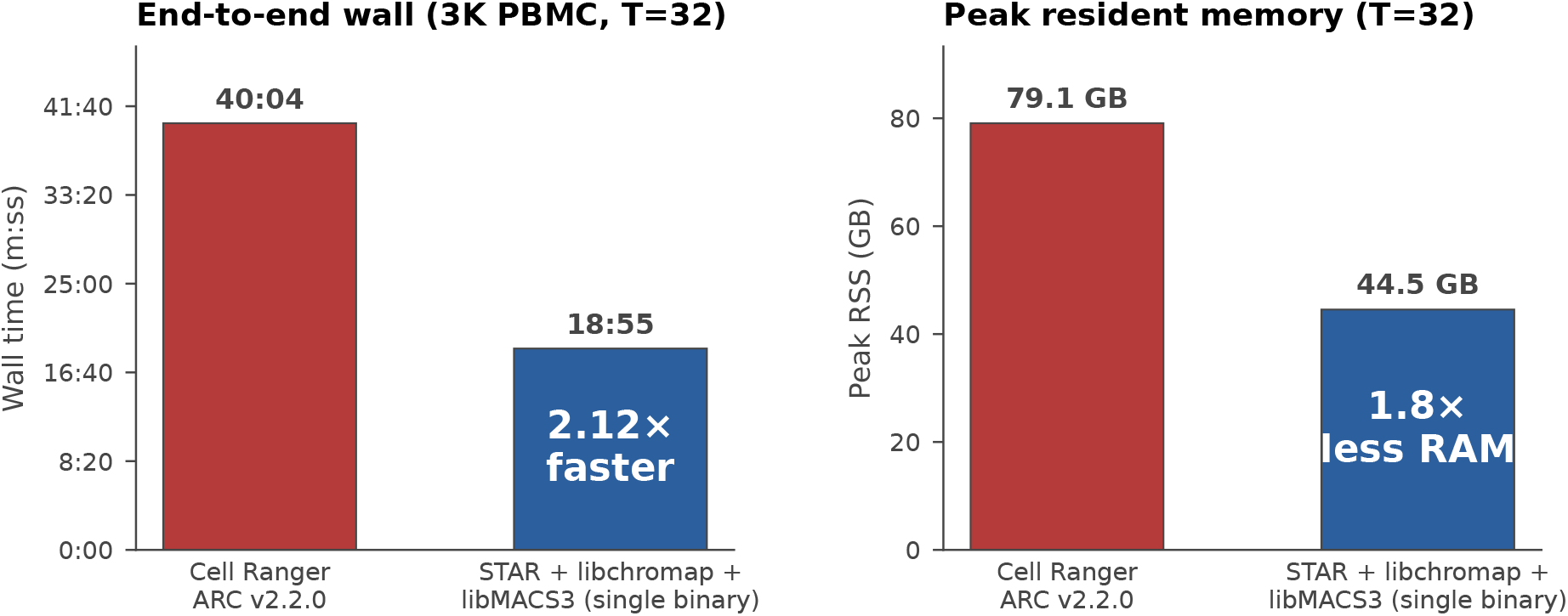
End-to-end multiomic benchmark on the public 10x 3K PBMC Multiome at 32 threads. Wall time (left) and peak resident memory (right) for Cell Ranger ARC v2.2.0 (red) versus the integrated single-binary STAR Suite + libchromap + libMACS3 pipeline (blue). The integrated pipeline completes in 18:55 wall and 44.55 GB peak RSS vs 40:04 and 79.07 GB for ARC—a 2.12× wall speedup with 1.8× less peak memory. Scope is matched: alignment, GEX UMI counting, GEX `EmptyDrops_CR`, ATAC mapping, narrow peak calling, and ATAC per-barcode cell call. ARC’s RSS is read from cellranger’s own `_perf` JSON because the parent-process `/usr/bin/time -v` reading is misleading (ARC forks 252 child processes at peak); the integrated pipeline’s RSS reading is single-process and accurate.

#### Low-memory spillover and concurrency rewrite

`libchromap`’s low-memory spillover path was rewritten as new architecture rather than patched. Per-thread overflow writers feeding a *k*-way merge on read-back replace the prior shared-buffer + atomic-write design, which had been deprecated as it could not safely combine with native BAM output, fragment generation, or in-process peak calling. The rewrite supports the cross-product of `--low-mem` with `--atac-fragments`, native BAM output, `--macs3-frag-low-mem`, and Y-filtering modes, and worker threads coordinate with the native BAM writer, the permit allocator (Section 5.1), and libMACS3 peak-call paths under the existing OpenMP scheduler. A pre-existing race condition in the legacy spillover code path that produced silent read drops at ~1*/*10^4^ on typical datasets and intermittent hangs at production scale (10^9^ reads) is resolved as a side effect of the rewrite (file references in `HISTORY.md`). The work was developed earlier for the MorPhiC consortium pipeline; see Section 5.3.

#### Y-chromosome filtering (utility)

A contig-name-pattern detector (`src/y_contig_detector.h`) routes Y-chromosome reads to optional separate output streams (`--emit-Y-bam`, `--emit-noY-bam`) for use cases that exclude or analyse Y separately. The detector runs at the output-writer layer, below the preset dispatch, and is therefore preset-agnostic in the implementation; current regression coverage exercises it on ATAC and on a generic paired-end synthetic reference (`tests/e2e/test_y_streams.sh`), and explicit cases for ChIP-seq and Hi-C presets are a planned extension to the matrix.

#### Regression suite covering Chromap’s existing presets

Chromap-suite ships with a regression matrix (cases C01–C11) that auto-validates the non-ATAC paths that Chromap Suite inherits unchanged from Chromap, alongside the ATAC paths the new work targets. The four assay presets the underlying Chromap supports—bulk ATAC (`--preset atac --BED`), single-cell ATAC (`--preset atac` + barcode whitelist), ChIP-seq (`--preset chip --TagAlign`), and Hi-C (`--preset hic` `--pairs`)—are exercised at S0 and S1 so that no addition described above introduces a regression in upstream Chromap behaviour. The S0 hermetic tier (`tests/run_libchromap_core_smoke.sh`) drives synthetic fixtures for each of the four presets plus the new utility flags (sorted BAM and index, low-memory BED parity, ATAC BAM + fragments, Y/noY split, MACS3 narrow peak calling) and runs in seconds for pre-commit gating. The S1 ENCODE tier (`tests/run_encode_cross_assay_smoke.sh`) runs the same four presets against downsampled real ENCODE accessions, with manifests committed in-tree and FASTQs cached out-of-tree. The S2 tier reserves the 100K and paper-fixture cases for heavier gates and is, in this version of the matrix, ATAC-focused; extending S2 to the full four-preset surface is planned. The regression suite is itself a substantive correctness contribution beyond the new functionality enumerated above: the upstream Chromap repository carried no comparable cross-preset test matrix, and we adopted three-tier testing under the same AI-assisted methodology described in Section 5.3. The S0 tier is mandatory for pre-commit checks; S1 and S2 are opt-in for hosts with appropriate fixture caches.

#### Agent and human accessibility

`libchromap` ships with a Model Context Protocol (MCP) server (`mcp_server/`) and a browser Launchpad UI that expose its workflows as schema-validated recipes. Ten chromap-specific workflow recipes (index build, BED and BAM-dual ATAC, ChIP TagAlign, Hi-C pairs, sorted BAM, Y/noY split, MACS3 narrow peak calling, ATAC evidence-from-peaks, and library-runner parity) double as both agent tool-call schemas (consumed via MCP-compatible clients) and human-facing recipe templates (rendered into command lines by the Launchpad UI). The MCP server exposes tools for workflow discovery, parameter validation, command rendering, and build management; a single set of YAMLs serves both audiences. The motivation is the same on both sides: replacing freeform shell-script orchestration with schema-validated workflow recipes reduces error-prone tool-boundary navigation, whether the operator is a human reading a recipe or an agent composing tool calls. The in-repository recipes are the per-binary surface; production-deployment recipes that orchestrate Chromap Suite alongside STAR Suite for the MorPhiC consortium’s multiomic releases live in a separate companion repository, `morphic-recipes` [11], and per-run reproducibility records (commit SHA, binary checksums, rendered command line, input/output inventories with checksums) are maintained in `morphic-provenance` [10].

### 5.2 libMACS3: a portable C++ narrow peak caller

MACS3 [15] is the de facto standard narrow peak caller for ATAC-seq and ChIP-seq analyses. It is implemented in Cython and compiles against the CPython runtime; embedding it in a non-Python binary therefore requires bringing in the full Python interpreter, which is impractical for a single-binary integrated pipeline. We reimplemented MACS3’s narrow peak-calling capability natively in C++ as `libMACS3`, a standalone library that exposes the capability through a clean C++ API and can be embedded directly in any C++ context (source: https://github.com/morphic-bio/libMACS3).

#### Validation

On the public 10x 3K PBMC Multiome ATAC channel (53.97 M fragments after deduplication), `libMACS3` produces output that is byte-identical to MACS3 v3.0.3 reference output across every artefact we validate. Byte identity is established by comparing the md5 sums of the full output files; per-artefact md5 values are recorded in the repository’s benchmark log and are not reproduced here. The artefacts checked are:

- narrowPeak file: 50,274 peaks;
- summits BED (default `--macs3-frag-summit-parity`);
- treat, lambda, and `ppois` bedGraphs (matching MACS3’s `-B` output);
- per-peak score columns of the narrowPeak file (signalValue, −log_10_(*p*), −log_10_(*q*)): agreement to within 10^*™*5^ float round-off.

#### Implementation methodology

Peak identification, statistical fields, and summit calling were re-implemented in C++ from MACS3’s published algorithmic specification, with output-based parity testing against MACS3 v3.0.3 reference outputs at each stage as the validation harness. One behaviour—the per-event tie-break in MACS3’s internal `pos_array` used during summit calling— required source-inspected adaptation under MACS3’s BSD-3 licence to reach byte-identical summits in degenerate multi-way-tie cases, and `libMACS3` is consequently distributed under BSD-3 rather than the MIT licence used elsewhere in Chromap Suite. A flag is available for users who prefer the documentation-faithful behaviour.

#### Performance

Standalone wall-clock and memory benchmarks on the same input (Fig. 1). Single-threaded, `libMACS3` completes in 92 s versus 721 s for MACS3 v3.0.3 (7.8× faster), with peak resident memory of 2.77 GB versus MACS3’s 2.50 GB (~10% overhead). At four threads, `libMACS3` completes in 70 s (10.3× faster than single-threaded MACS3) at the same 2.77 GB peak memory: memory parity at the practical thread budget. At 24 threads, `libMACS3` reaches 64 s (11.3×) at 3.88 GB; speedup scales sublinearly beyond four threads because the per-chromosome serial sweep dominates the remaining wall time. All thread counts produce byte-identical narrowPeak.

#### API and scope

`libMACS3` exposes the narrow peak-calling capability through a single public entry point, `RunMacs3FragPeakPipelineFromSortedIterator`, which consumes a fragment iterator and produces narrowPeak and summit outputs. The fragment-iterator abstraction allows callers to feed fragments from a TSV file (`OpenFileFragmentIterator`) or directly from memory (`WrapVectorFragmentIterator`). Standalone `libchromap` drives the in-memory path; the STAR Suite multiomic pipeline drives libMACS3 through the AEV1 binary sidecar, consumed by `RunMultiomeAtacPeakMex` as an in-process post-alignment phase inside STAR (Section 2.3). Scope is deliberately narrow: only narrow peak calling is provided. MACS3’s other operations— HMM-based broad peak calling, `refinepeak`, and `bdgcmp` as standalone paths—are out of scope. `libMACS3` is a capability port, not an API replacement for Cython MACS3.

### 5.3 AI-assisted implementation methodology

#### Scope

The new components built for this work—libMACS3, the extraction of libchromap as a callable library, and the multiomic integration into STAR Suite—were developed using AI-assisted programming under explicit human direction. We separate this scope from the pre-existing infrastructure on which the result depends: Chromap’s low-memory spillover rewrite, native BAM output, and STAR Suite’s permit allocator (Section 5.1) were developed earlier for the MorPhiC consortium pipeline, and STAR Suite’s underlying scheduler infrastructure substantially pre-dates AI-assisted development. The methodology notes below apply to the new components only.

#### libMACS3: meta-prompted decomposition before generation

End-to-end “clean-room generation” of a non-trivial algorithm from documentation alone produced poor parity in our hands (initial Jaccard 0.2–0.3 against MACS3 reference output, unguided) and is not a workable pattern. The pattern that worked was meta-prompted decomposition: rather than asking the model to implement the algorithm directly, we asked it first to assess whether MACS3’s narrow peak-calling could be broken down into discrete stages amenable to per-stage parity testing, and to proceed stage-by-stage if so. The model produced the decomposition (pileup, peak boundary detection, summit calling, statistical scoring) and the per-stage parity tests against MACS3 reference outputs; once the decomposition and harness were in place, the clean-room implementation against documented specification reached approximately 98% summit parity (and full byte-identity on most output artefacts) in essentially one implementation pass, with one remaining tie-break edge case at the summit-calling stage that required source-inspected adaptation (next paragraph) before byte-identical narrowPeak and summits emerged. The human contribution at this step was the meta-prompt itself, not the algorithmic decomposition. Subsequent human-directed optimization (binary search in place of linear search, per-chromosome parallelisation, sweep-line workspace) brought the standalone runtime to the 7.8–11.3× speedup over Cython MACS3 reported in Section 5.2.

#### Summit edge cases under permissive licensing

Summit calling reached approximately 98% within 1 bp of the MACS3 reference through clean-room implementation against documented specification, but the remaining ~2% showed deviations of up to ~222 bp in degenerate cases (multi-way ties of identical pileup-position scores), despite faithful implementation of the documented tie-breaking rule. Subsequent inspection of MACS3’s Cython source—permitted under MACS3’s BSD-3 licence—revealed that the actual summit tie-resolution had diverged from public documentation (MACS3’s internal `pos array` is per-event rather than per-distinct-value as documented). Reproducing the actual algorithm restored byte-identity (Section 5.2). Permissive licensing was therefore load-bearing: a more restrictive licence on the original tool would have left libMACS3 stuck at the 98% asymptote.

#### libchromap extraction and STAR Suite integration

Extracting libchromap as a callable library was largely mechanical refactoring under human direction: the existing `chromap` CLI binary was retargeted to call `libchromap.a` through the `libchromap_contract` API rather than embedding the mapping logic directly. Wiring `libchromap_contract` into STAR Suite took approximately half a day: installing forwarding shims, wiring the configuration paths, and unifying htslib so a single library instance handles both GEX and ATAC BAM output. The allocator-side work was more substantive. STAR Suite’s permit allocator originally targeted two concurrent processing domains (GEX mapping and feature processing); supporting ATAC as a third domain required extending the underlying wake-up and scheduling logic and evaluating several thread-distribution strategies before settling on the one that consistently gave the shortest wall time (Section 5.1). We took the opportunity to generalise the allocator so that registering further concurrent domains no longer requires per-domain code changes; this extension is what makes the headline 18:55 wall reproducible and what positions the allocator to support additional modalities without further redesign.

## Declarations

### Ethics approval and consent to participate

Not applicable. The 3K PBMC Multiome benchmark data is publicly available from 10x Genomics; no new human or animal data were generated.

### Consent for publication

Not applicable.

### Availability of data and materials

The source code described in this work is freely available under permissive open-source licences:

- **libMACS3** (BSD-3): https://github.com/morphic-bio/libMACS3
- **Chromap Suite / libchromap** (MIT): https://github.com/morphic-bio/Chromap-suite
- **STAR Suite** (MIT, with multiomic integration via libchromap + libMACS3): https://github.com/morphic-bio/STAR-suite
- **morphic-recipes** [11] — production-deployment recipes for Chromap Suite and STAR Suite as used by the MorPhiC consortium: https://github.com/morphic-bio/morphic-recipes
- **morphic-provenance** [10] — per-run reproducibility records (commit SHA, binary checksums, rendered command line, input/output inventories with checksums) for MorPhiC releases: https://github.com/morphic-bio/morphic-provenance

The 10x Genomics 3K PBMC Multiome dataset used for benchmarking is publicly available from 10x Genomics [1]. Full reproduction artefacts for the figures and benchmarks in this paper—including benchmark TSVs and figure-render scripts—are included in the paper repository alongside this manuscript.

### Authors’ contributions

L.H.H. conceived the project, implemented the software, conducted the benchmarks, and drafted the manuscript. K.Y.Y. supervised the work and revised the manuscript. Both authors approved the final manuscript.

## Acknowledgments

We thank Bill Flynn at the Jackson Laboratory for feedback on the multiomic approach, and Jesse Engreitz, Anshul Kundaje, and Amanda Everitt at Stanford University for feedback on ATAC-seq methodology.

We acknowledge the authors of Chromap (Haowen Zhang and colleagues) and MACS3 (Tao Liu and colleagues) for the original tools that motivated this work; libchromap is a derivative of Chromap distributed under its MIT license, and libMACS3 is a clean-room implementation of MACS3’s narrow peak-calling capability with source-inspected adaptation for summit edge cases distributed under MACS3’s BSD-3 license.

